# Swimming motions evoke Ca^2+^ events in vascular endothelial cells of larval zebrafish via mechanical activation of Piezo1

**DOI:** 10.1101/2025.02.05.636757

**Authors:** Bill Z. Jia, Xin Tang, Marlies P. Rossmann, Leonard I. Zon, Florian Engert, Adam E. Cohen

## Abstract

Calcium signaling in blood vessels regulates their growth^1,2^, immune response^3^, and vascular tone^4^. Vascular endothelial cells are known to be mechanosensitive^5–7^, and it has been assumed that this mechanosensation mediates calcium responses to pulsatile blood flow^8–10^. Here we show that in larval zebrafish, the dominant trigger for vascular endothelial Ca^2+^ events comes from body motion, not heartbeat-driven blood flow. Through a series of pharmacological and mechanical perturbations, we showed that body motion is necessary and sufficient to induce endothelial Ca^2+^ events, while neither neural activity nor blood circulation is either necessary or sufficient. Knockout and temporally restricted knockdown of *piezo1* eliminated the motion-induced Ca^2+^ events. Our results demonstrate that swimming-induced tissue motion is an important driver of endothelial Ca^2+^ dynamics in larval zebrafish.

**Highlights:**

- Swimming motions in larval zebrafish evoke large, rapid, and pervasive Ca^2+^ transients in vascular endothelial cells.
- These Ca^2+^ transients do not require neural firing, muscular electrical activity, endocrine factors, or heart-driven blood flow.
- Mechanical forces are necessary and sufficient for endothelial Ca^2+^ transients.
- Endothelial Ca^2+^ transients require the mechanosensitive ion channel Piezo1.

## Introduction

Vascular endothelial cells (ECs) experience a range of distinct mechanical forces *in vivo*^1,2,6,7^. These include shear stress from blood flow, radial stretch from blood pressure, circumferential stretch from smooth muscle-mediated vasodilation, and, in some parts of the animal, axial stretch from skeletal muscle-mediated body motion^6,11^ (Fig. S1). Each has a characteristic magnitude and timescale (Supplementary Table 1). In principle, these different modes of stimulation could activate distinct signaling pathways and cellular responses^12–14^. Mechanical perturbation experiments on cultured cells or explants typically impose stresses that differ in magnitude and direction from the forces encountered *in vivo*^5,15,16^, and may miss aspects of the biochemical environment^1^. Thus *in vitro* or *ex vivo* experiments cannot readily assign biochemical responses to specific sources of mechanical stress *in vivo*.

Endothelial cells show Ca^2+^ events under a variety of conditions, including activation of shear stress^9^, variation of membrane potential^17^, stimulation by ATP^18^ and exposure to VEGF^19^ or inflammatory mediators^20^. Though many experiments have probed Ca^2+^ signaling due to flow-induced shear stress in cultured cells and explants^21,22^, experiments *in vivo* are required to establish the natural triggers of EC Ca^2+^ signaling. Experiments in zebrafish found that early in development, flow-induced deflection of EC cilia led to Ca^2+^ influx^9^. Flow-induced Ca^2+^ events, however, were only observed early in embryonic development (< 2 dpf (days post fertilization) )^9^. Whether baseline blood flow is important in triggering EC Ca^2+^ impulses *in vivo* at later developmental stages remains unclear.

Increased blood flow and swelling of active skeletal muscles (“muscular hyperemia”) is a familiar experience to anyone who has lifted weights, though the mechanisms underlying this effect are not fully established^23^. Muscular activity in the hamster cremaster muscle induces EC Ca^2+^ events *in vivo*, which in turn induce arteriolar vasodilation^24^. Thus EC Ca^2+^ dynamics play a crucial role in regulating changes in blood flow in response to metabolic demand^25,26^. However, the molecular mechanisms by which the muscular activity induces the EC Ca^2+^ events have remained unclear^27^. Hypotheses include sympathetic neuro-vascular coupling^28^, release of neurotransmitter (e.g. acetylcholine)^29^, changes in oxygen availability^30^ or changes in metabolite concentration^31,32^. In contrast to well-studied shear force activation, relatively few studies on direct mechanical activation of EC Ca^2+^ dynamics by skeletal muscle contraction have been performed *in vivo*. When one considers quantitatively the magnitudes of the EC deformations that arise under different physiological forces (Supplementary Table 1), motion of skeletal muscle emerges as the largest and fastest natural mechanical input. Thus, it is natural to ask what role body motion plays in inducing EC Ca^2+^ events *in vivo*, and what mechano-sensitive molecules are responsible for mediating these responses.

We studied EC Ca^2+^ dynamics in embryonic zebrafish. Due to its small size, rapid development, genetic tractability, and optical transparency, the zebrafish is an excellent model for studying endothelial signaling *in vivo*^33,34^. In zebrafish expressing a genetically encoded fluorescent Ca^2+^ indicator in vascular endothelial cells, we observed dramatic calcium transients associated with body motion. We then used pharmacological, mechanical, and genetic perturbations to explore the coupling between body motion and EC Ca^2+^ events. These experiments established that the interaction was through direct mechanical coupling, mediated by the mechano-sensitive ion channel, Piezo1.

## Results

### Body motion of larval zebrafish triggers Ca^2+^ events in endothelial cells *in vivo*

We performed whole body fluorescence imaging of transgenic larval zebrafish expressing the GCaMP5G Ca^2+^ indicator under control of the EC-specific *kdrl* promoter *Tg*(*kdrl: Gal4: UAS: GCaMP5g*) (Fig. 1A). Resting fish showed few spontaneous EC Ca^2+^ events. However, when the fish performed an escape response tail flick, dramatic Ca^2+^ events arose in most dorsal aorta (DA), caudal artery (CA), intersegmental vessels (ISVs), and dorsal longitudinal anastomotic vessels (DLAV; Fig. 1B, Supplementary Video 1), but not in dorsal longitudinal vein (DLV) vessels in brain, the caudal vein (CV), or posterior cardinal vein (PCV) vessels in trunk. In intersegmental vessels, the median onset time of Ca^2+^ transients was 0.44 (0.04 – 1.94) s after the start of an escape twitch and the transients decayed with a median time constant of 22.5 (19.8 – 26.6) s (median (IQR), *n* = 31 events in two 5 dpf fish; Fig. S2). Motion-induced Ca^2+^ events were observed from 3 to 7 dpf. A transgenic zebrafish line expressing an eGFP marker in the ECs, *Tg (kdrl: Gal4: UAS: eGFP)*, did not show motion-induced fluorescence transients, confirming that the signals observed in the GCaMP5g fish were due to changes in brightness of the Ca^2+^ indicator, and not due to motion artifacts (Fig. S3).

**Figure 1:**
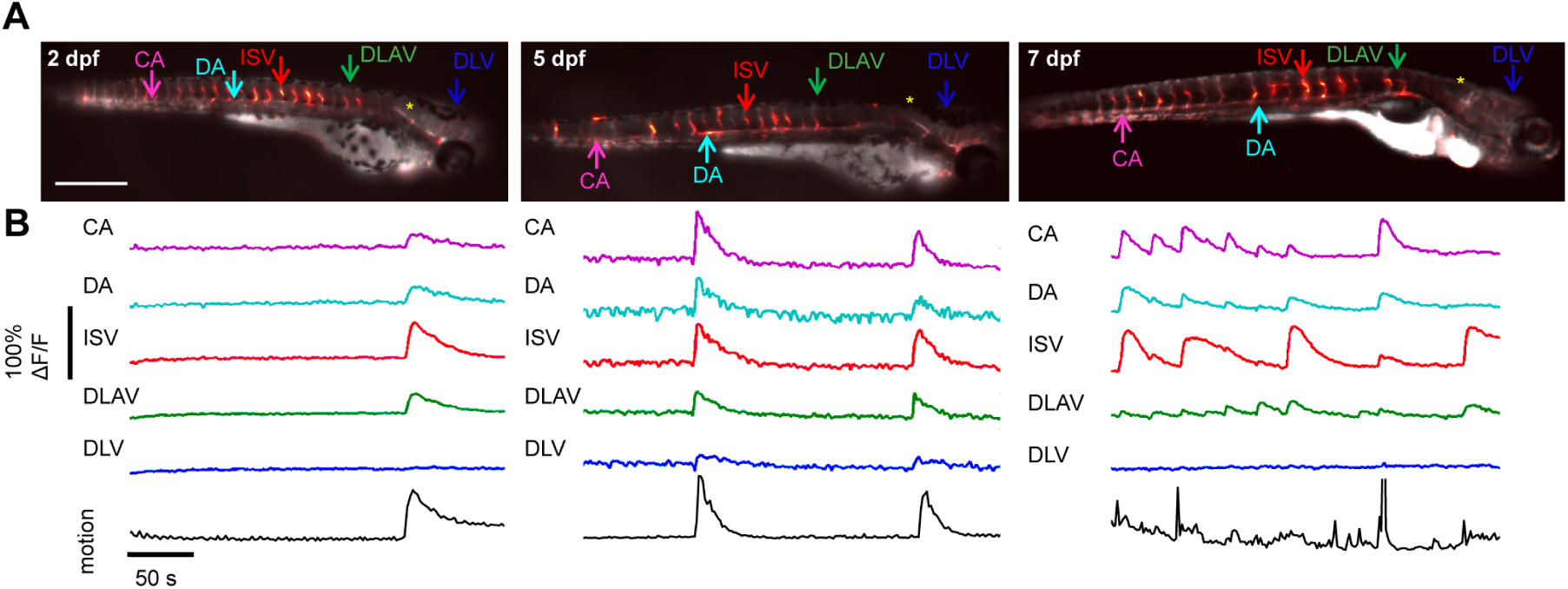
Vascular endothelial cell calcium events are associated with spontaneous swim motions in larval zebrafish. **(A)** Vascular ECs of larval zebrafish show Ca^2+^ events at 2, 5, and 7 dpf. Images are a composite of basal fluorescence (greyscale) and maximum ΔF/F (heatmap). Scale bar: 500 μm. **(B)** ΔF/F time-traces of Ca^2+^ dynamics on dorsal aorta (DA), caudal artery (CA), intersegmental vessels (ISVs), dorsal longitudinal anastomotic vessels (DLAV), and dorsal longitudinal vein (DLV) ^31,32^. Calcium events were not observed in hindbrain vessels (indicated by yellow asterisk “*”). Black solid lines indicate spontaneous motion.

A previous report described elevation in basal Ca^2+^ levels in embryonic zebrafish ECs, induced by hemodynamic deflection of primary cilia^9^. That effect did not include Ca^2+^ dynamics on the timescale of seconds and disappeared by 48 hours post-fertilization (hpf) due to disassembly of cilia^9,35^. The body motion-induced Ca^2+^ transients described here appear to be a distinct phenomenon.

### Skeletal muscle contraction directly triggers EC Ca^2+^ transients

We next sought to identify the cellular and molecular pathways that led to these motion-associated EC Ca^2+^ transients. We first examined the necessity and sufficiency of different physiological dynamics involved in zebrafish swimming motion: (1) motor circuit neural activity; (2) cardiac function; and (3) skeletal muscle contraction.

We exploited the aversive visual-motor reflex of zebrafish larvae to violet light^36^ to trigger a reproducible and naturalistic series of escape twitches leading to EC Ca^2+^ activation (Fig. 2A; Fig. S4). We simultaneously imaged Ca^2+^ activity in ECs and in glutamatergic excitatory interneurons of the spinal cord using *Tg(kdrl:Gal4;vglut2a:Gal4;UAS:GCaMP6s)* fish^37^. Spinal glutamatergic interneurons synapse onto cholinergic motor neurons and are a common proxy for neural motor activation in larval zebrafish^38,39^. We focused on intersegmental ECs to sample within each field of view multiple distinct vessels that had similar geometries and experienced similar mechanical forces. Violet light (405 nm) illumination of the eyes in 5 dpf larvae for 10 s elicited robust Ca^2+^ activity in spinal glutamatergic neurons (98% of trials), twitch responses (98% of trials) and EC Ca^2+^ events (89% of trials; *n* = 9 fish, 54 total trials; Figs. 2B – D; Supplementary Video 2).

**Figure 2:**
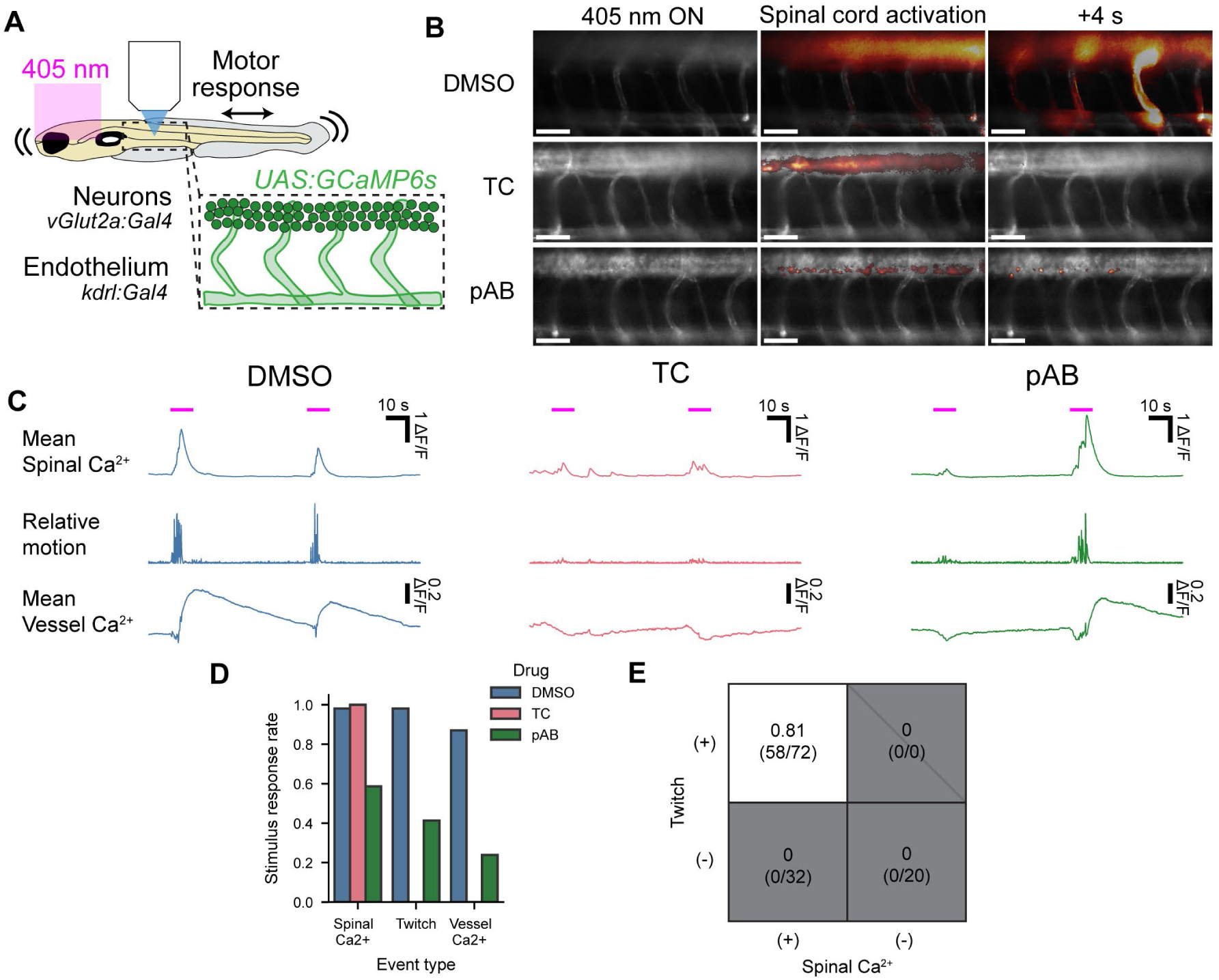
Calcium events in ECs are triggered by muscle contraction and not neural activity or cardiac output. **(A)** Experimental setup. Violet light on the eyes evoked escape attempts; Ca^2+^ responses were monitored in the spinal cord and in vascular endothelial cells. **(B)** Typical response profiles in response to vehicle control (DMSO), tubocurarine (TC), or para-amino blebbistatin (pAB). Scale bars 50 µm. **(C)** Typical response waveforms in response to DMSO, TC, or pAB. The negative-going transients in the vessel Ca^2+^ signals are motion artifacts. **(D)** Quantification of response probabilities by different measures in response to DMSO, TC, or pAB. **(E)** Probability of EC Ca^2+^ transients under different conditions of twitch and spinal Ca^2+^ responses over all treatments. Events with spinal Ca^2+^ but no twitch did not evoke EC Ca^2+^ transients, while events with twitch did evoke EC Ca^2+^ transients with 81% probability. *n* = 9 fish DMSO, 4 fish TC, 8 fish pAB.

To assess whether neural activity alone could evoke EC Ca^2+^ events, we applied tubocurarine (TC, 2.2 mM), which blocks nicotinic acetylcholine receptors (nAchRs) and thus disrupts signal transmission across neuromuscular junctions. In the presence of TC, we still observed neural Ca^2+^ responses to 100% of stimuli (*n* = 4 fish, 24 total trials) but we detected neither physical twitch responses nor intersegmental EC Ca^2+^ responses (Fig. 2B – D; Supplementary Video 2). We observed stimulus-correlated Ca^2+^ transients localized to the junction between intersegmental arteries and the dorsal aorta (11/24 trials; Supplementary Video 3). These events were distinct in magnitude, timing, and location compared to the motion-induced EC Ca^2+^ transients. We speculate they may have been caused by neurovascular coupling or changes in flow due to cardiac output. Due to their clear differences from the motion-evoked events, we excluded them from analysis.

In TC-treated fish, the amplitude of the neural Ca^2+^ response was smaller, on average, than in drug-free controls, perhaps because of absence of somatosensory feedback (Fig. S4). Nonetheless, comparing events of matched spinal Ca^2+^ amplitude, the EC Ca^2+^ response remained significantly smaller in the TC-treated fish than in the drug-free controls (*p* = 1.3e-4, mixed-effects general linear model; Fig. S4). Together, these observations show that spinal motor circuit activity alone was insufficient to evoke EC Ca^2+^ transients.

A further possibility is that electrical excitation of skeletal muscles somehow activated EC Ca^2+^ transients, even in the absence of motion. To rule out this possibility, we applied para-amino blebbistatin (pAB, 50 µM), a photostable myosin II inhibitor which blocks muscle contraction^40^ without impairing muscle electrical excitation. This drug partially suppressed spinal Ca^2+^ responses (responses observed in 27/46 trials, *n* = 8 fish), and was only partially effective at blocking twitches (twitches observed in 19/46 trials, all accompanied by spinal Ca^2+^ responses; Fig. 2B – D; Supplementary Video 2). This partial efficacy allowed observation of distinct combinations of spinal activity and twitching. EC Ca^2+^ responses occurred in 11/19 of the trials which evoked twitches, and not in any of the 8 trials which evoked spinal Ca^2+^ activity but no twitches nor in any of the 19 trials which evoked no spinal Ca^2+^ activity. These results confirm the necessity of twitches for EC Ca^2+^ responses (Fig. 2E; Fig. S4). Suppression of EC Ca^2+^ by pAB also establishes that the previously observed suppression by TC was not mediated through direct inhibition of endothelial nAchRs^41^. pAB also severely disrupted the heartbeat, eliminating resting blood flow (Supplementary Video 4). Persistence of the EC Ca^2+^ events in the absence of a heartbeat implies that neither circulating endocrine factors nor changes in vessel mechanical forces driven by cardiac output were required for motion-induced EC Ca^2+^ events.

### Mechanical stimuli induce EC Ca^2+^ transients via mechanosensitive ion channels

Next, we sought to determine whether EC Ca^2+^ events were driven by the mechanical forces exerted by muscle contraction, or by some other downstream physiological process We set up a closed-loop mechanical pusher system to apply controlled mechanical stimuli with physiologically relevant magnitude^42^ to the whole body of larval zebrafish (Fig. 3A). In the presence of TC, application or release of a 5-gram weight distributed over the fish induced EC Ca^2+^ events in 6 of 9 fish ranging from 3-7 dpf. As before, TC completely suppressed twitch motion (Fig. 3B, C). These findings imply that mechanical forces alone are sufficient to induce EC Ca^2+^ events.

**Figure 3:**
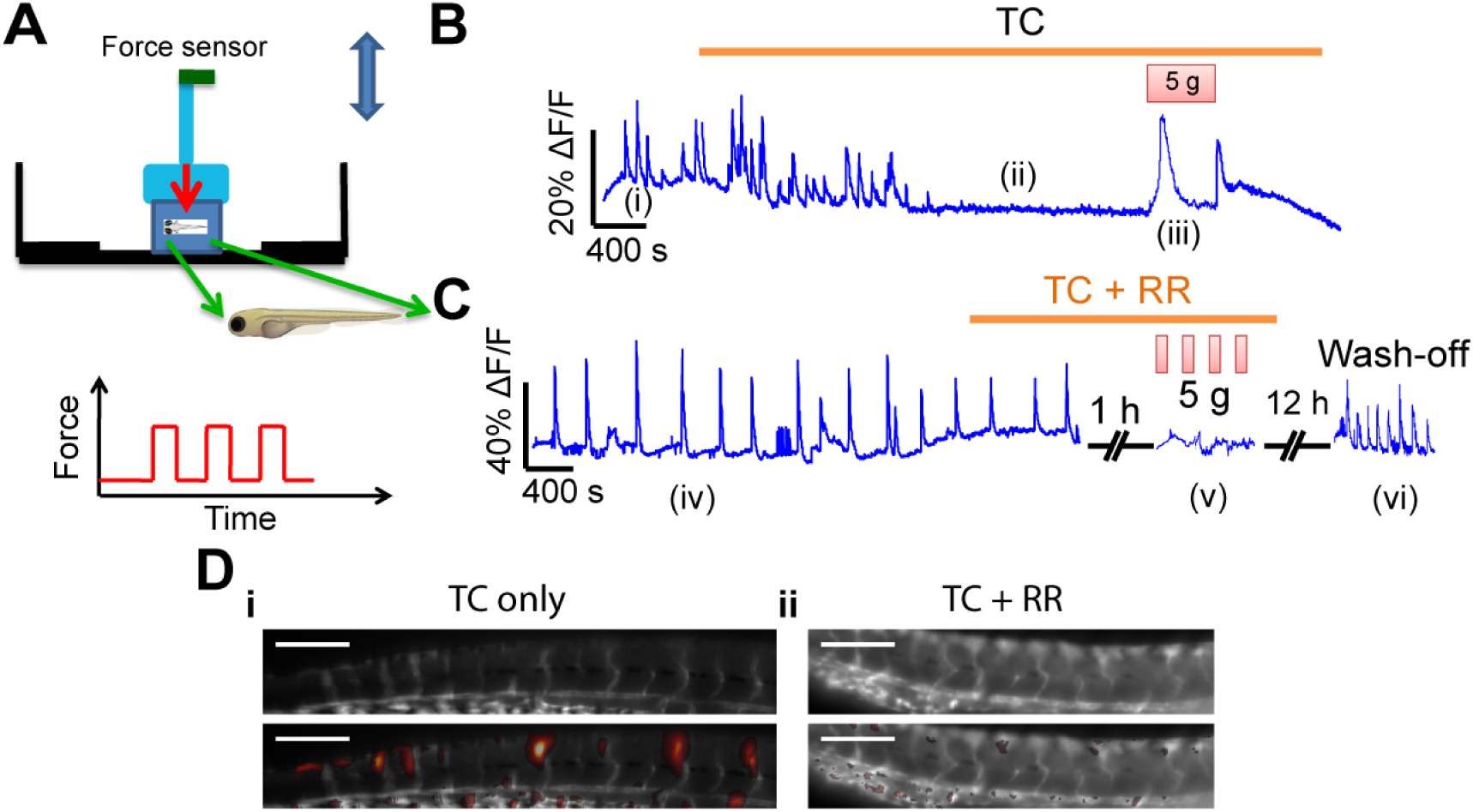
Mechanical stimuli induce Ca^2+^ events in ECs. **(A)** Schematic of closed-loop mechano-pusher. **(B)** Calcium dynamics of ECs in zebrafish under consecutive conditions: (i) spontaneous twitching, (ii) blockage of neuromuscular junction by tubocurarine (TC), and (iii) stimulation with mechanical force. **(C)** Calcium dynamics of ruthenium red (RR) treated ECs under consecutive conditions: (iv) spontaneous twitching, (v) mechanical stimulation with TC and RR, and (vi) drug wash-out. **(D)** Examples of changes in EC GCaMP5G intensity from rest (top) after mechanical push (bottom). Scale bars 200 µm.

Based on the rapid response of EC Ca^2+^ to spontaneous motion and the sufficiency of mechanical deformation to trigger EC Ca^2+^ events, we hypothesized that mechano-sensitive ion channels on ECs might mediate entry of extracellular Ca^2+^ ions. We therefore examined EC Ca^2+^ events in zebrafish treated with ruthenium red (RR, 50 μM), a non-specific blocker of stretch-activated cation channels, including Piezo and TRP family channels^43,44^. To isolate purely mechanically driven Ca^2+^ events, we paralyzed larvae with TC. Addition of RR to the curare-containing incubation medium abolished the mechanically induced EC Ca^2+^ events (Fig. 3C, D, 0/3 animals with EC Ca^2+^ events). After RR and TC wash-out, zebrafish produced spontaneous body motions and associated EC Ca^2+^ events (Fig. 3C), confirming that the effect of the drugs was reversible. These results suggest that mechano-sensitive ion channels, possibly in the EC or neighboring coupled cells, are required for the mechano-sensation process in EC calcium signaling.

### Piezo1 is required for mechano-activated EC Ca^2+^ events

To determine the molecular identity of the ion channels involved in EC mechanotransduction, we turned to a CRISPR/Cas9-mediated knockout (KO) strategy. We selected three candidates based on Ca^2+^ permeability, sub-second kinetics^45–47^, and previously documented expression in endothelial cells^46,48,49^: Piezo1, PKD2, and TRPV4. We tested each KO by measuring EC Ca^2+^ events during electric field stimulus-evoked body motions in live fish.

Single guide RNAs (sgRNAs) targeting *piezo1* diminished protein expression as quantified by Western blot (Fig. 4A), and successful editing of *piezo1* was confirmed by gel electrophoresis and sequencing (Fig. S5). In *Tg (kdrl:Gal4; UAS:GCaMP5G)* fish with *piezo1* KO, we observed significantly diminished motion-triggered EC Ca^2+^ events (Fig. 4B). In contrast, uninjected fish showed normal motion-triggered EC Ca^2+^ events (Fig. 4B – D). *pkd2* and *trpv4* KO fish had no detectable difference in motion-triggered EC Ca^2+^ events compared to uninjected controls (Fig. 4 B – D).

**Figure 4:**
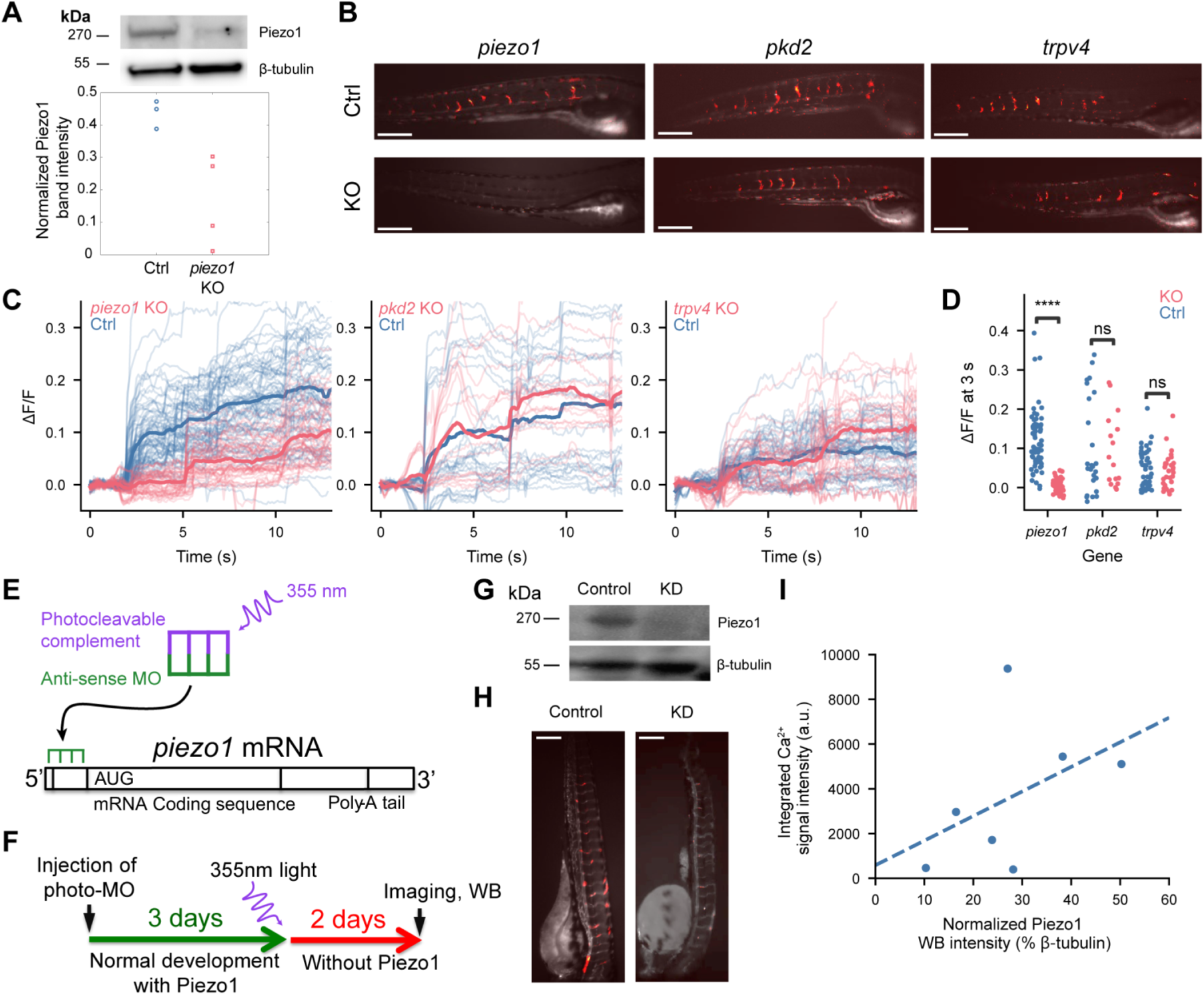
Piezo1 channels are required for motion-evoked EC Ca^2+^ events. **(A)** western blot quantification of Piezo1 expression in whole-animal lysates from control and Piezo1 gRNA-injected fish (5 dpf), normalized to β-tubulin band intensity. **(B)** Relative GCaMP5G fluorescence 3 s after electrical field stimulus in knockout (*piezo1*, *pkd2*, or *trpv4*) versus control *Tg(kdrl:Gal4;UAS:GCaMP5G)* fish. Scale bars 500 µm. **(C)** ΔF/F traces for individual blood vessel segments. Electrical field stimuli at 2, 6, and 11 s. Solid lines indicate mean over all measured vessels in each group of fish. **(D)** ΔF/F 3 seconds after the first electrical field stimulus. (C – D) *n* = 6 controls, *n* = 3 KO *piezo1*; *n* = 3 controls, *n* = 3 KO *pkd2*; *n* = 4 controls, *n* = 3 KO *trpv4*. **(E)** Photo-morpholino (MO) mechanism of action. **(F)** Experimental design for temporally restricted photo-morpholino knockdown of Piezo1. **(G)** Western blot measuring Piezo1 knockdown by photo-MO. (H) Relative GCaMP5G fluorescence 3 s after electrical field stimulus in scrambled and *piezo1* photo-MO treated fish. **(I)** Integrated calcium signal intensity over whole fish 3 s after electrical field stimulus compared to post-hoc western blot Piezo1 band intensity normalized to β-tubulin band intensity (*R^2^*= 0.20). Statistical tests: (D) Mixed-effects linear model considering individual fish and CRISPR/Cas9 treatment of each vessel, *piezo1*: *z* = -4.827, *p* = 1.38e-6; *pkd2*: *z* = 0.481, *p* = 0.630; *trpv4*: *z* = 0.002, *p* = 0.998.

Endothelium-specific Piezo1 deficiency results in defects in vascular development and embryonic lethality: Piezo1^-/-^ mice embryos die mid-gestation at E9.5^10,50^. To control for possible developmental defects resulting from chronic Piezo1 deficiency, we selectively knocked down *piezo1* in zebrafish at 3 dpf, using a photo-cleavable translation-blocking morpholino (photo-MO; Fig. 4E). Embryos were injected with the photo-MO at the single-cell stage and exposed to 355 nm laser at 3 dpf (Fig. 4F). Western blot analysis confirmed Piezo1 knockdown (KD) in treated fish (Fig. 4G). All *piezo1* morphants showed normal vascular morphology with no noticeable development defects. By 5 dpf, we found that *piezo1* morphants showed reduced EC Ca^2+^ events following body twitching (Fig. 4H). We found a positive correlation between EC Ca^2+^ transient amplitudes and Piezo1 protein levels measured in the same fish (*R^2^* = 0.20, Fig. 4I). These data confirm that Piezo1 mediates rapid mechanically induced Ca^2+^ events in ECs of zebrafish *in vivo*.

## Discussion

We demonstrated a direct mechanical coupling between physical tissue motion and EC Ca^2+^ signaling *in vivo*, mediated by the Piezo1 mechanosensitive ion channel. Basal blood flow alone was not sufficient to trigger these Ca^2+^ events, while swimming-induced motions or externally applied compression induced these events even in the absence of cardiac function.

Mechanotransduction in vasculature has been implicated in regulating vascular tone, primarily through sensing changes in blood flow directly or consequent changes in oxygenation ^51,52^. Mechanosensitive responses to transient systemic changes, such as those induced by exercise, are often proposed to be downstream of changes in heart rate which in turn affect the blood pressure drop, flow rate, and shear stress in vessels^23,53^. On the other hand, changes in vascular tone have been observed as rapidly as a single muscular contraction (∼ 1 s or less)^54–59^, and in response to externally applied forces, leading some to propose that local mechanical compression of vessels may also play a role^59,60^. Here we show that EC compression by muscle action or external force is directly sensed by Piezo1, providing a molecular mechanism of triggering rapid cellular responses to local mechanical forces.

The overall function of Piezo1 in vascular physiology remains uncertain, and it may serve multiple competing functions. Ca^2+^ entry in endothelial cells induces vasodilatory responses by activating synthesis nitric oxide^61^, and through hyperpolarization via calcium-activated potassium channels^62,63^. On the other hand, Piezo1 activation directly provides a depolarizing cation current which may counteract endothelial hyperpolarization-induced vascular smooth muscle cell (VSMC) relaxation^63–65^.

Simultaneous imaging of Ca^2+^ and membrane voltage in zebrafish^66,67^ may allow direct dissection of the roles of Piezo1 in depolarization-driven vasoconstriction and in Ca^2+^-driven vasodilation. Optogenetic stimulation of endothelial cells, paired with Ca^2+^ ^68^ or voltage^69^ imaging could also help dissect these mechanisms. The magnitude or timescale of Piezo1 activation may differ between vessels and depend on context, and thus play a role in recruiting different cellular Ca^2+^-handling mechanisms or distinct Ca^2+^-sensing pathways^70^.

In mice, Piezo1 contributes to higher systemic blood pressure during exercise and to vasoconstriction in mesenteric resistance arteries (when NO synthesis is blocked)^64^. Similarly, Piezo1-mediated depolarization serves as a brake on activity-induced hyperemia in the brain vasculature^65^. In contrast, arteriolar dilation upon hamster cremaster muscle contraction requires EC Ca^2+^ elevation^24^. We did not observe significant modulation of vascular tone in our experiments. While larval zebrafish vessels at 5 dpf are responsive to α-adrenergic agonists and nitric oxide generators^71^, it is possible that mechanisms sensitive to Ca^2+^ or V_mem_ are not yet active at this age. Alternatively, Piezo1-mediated effects may not contribute to compression-mediated vascular tone modulation. Compression-induced vasodilation has been reported even in EC-denuded vessels^59^, indicating the existence of EC-independent mechanosensitive pathways for modulating vascular tone.

In embryonic development, forces related to blood flow are important for angiogenesis^9,10,50^, cardiac valve and endocardium development^72^, VSMC differentiation and arterial vasculature stabilization^73,74^, and hematopoietic stem cell induction and expansion^75,76^. Piezo1 is implicated in the transduction of these forces^10,50,74,76^. Our findings suggest that Piezo1 also integrates information on body motion that emerges during development. At a longer timescale, endothelial Piezo1 might also sense local forces exerted by neighboring tissues during morphogenesis^77^. Signaling dynamics induced by these local forces may also be medically relevant. For instance, recent studies suggest Piezo1 and other mechanosensitive channels in the endothelium have protective or pathological roles in ventilation-induced lung injuries^78–80^, although their roles in normal breathing are less certain^81^. Other pathologies where external forces may affect endothelial cells include pressure ulcers (“bed sores”), muscular atrophy, and tumor growth. Our work highlights the importance of mapping molecular mechanisms of mechanosensation in the context of physiological body forces.

## Materials and methods

### Zebrafish breeding and DNA constructs

Animal experiment protocols were approved by the Harvard University Institutional Animal Care and Use Committee (IACUC, protocol 10-13-4). All wild-type (WT) and transgenic zebrafish (*Danio rerio*) were raised and bred on a 14/10-hour light/dark cycle at 28.5 °C following standard procedures.

To obtain transgenic strains which express GCaMP5G in cytoplasm of vascular endothelial cells, the transgenic lines *Tg(kdrl:Gal4)*^82^ and *Tg(UAS:GCaMP5G)*^83^ was crossed, and offspring were screened by fluorescence expression pattern for imaging experiments at desired stages. To obtain larvae expressing GCaMP6s in vascular endothelial cells and glutamatergic interneurons, *Tg(kdrl:Gal4)* fish were crossed with *Tg(vglut2a:Gal4;UAS:GCaMP6s)* fish^37,84^.

### Microscopy and imaging

All embryos and larvae were raised in E3 media (5 mM NaCl, 0.17 mM KCl, 0.33 mM CaCl_2_, and 0.33 mM MgSO_4_) containing 0.003% 1-phenyl-2-thioura (PTU) to prevent melanogenesis during embryogenesis. Before imaging experiments, embryos and larvae were manually mounted in 2% low-melting-point agarose lateral-side down on glass-bottom petri-dish (MatTek Co.) for inverted microscope imaging, or on an agarose trench for upright microscope imaging.

To document expression pattern and morphology, fish were imaged on a Leica stereomicroscope, an Olympus dissection microscope (BX51W epifluorescence), or a custom-built ultra-widefield microscope. Functional calcium imaging, mechanical force application, violet light stimulation, and electrical field stimulation experiments were conducted for fish at 2-7 days post fertilization (dpf) using a custom-built epifluorescence microscope at room temperature. Illumination was provided by a solid state 488 nm laser (Coherent Obis 1226419, 100 mW), or a 488 nm 300 mW LED (LED Engin) driven by a current driver (Thorlabs DC4100).

Imaging was performed using 4x (Olympus Fluor, NA 0.24), 10x (Olympus UPlanSApo, NA 0.40; XLPLN10XSVMP, NA 1.0) objectives. A quad-band emission filter (Semrock Di01-R405/488/561/635) separated fluorescence from excitation light. Fluorescence passed through a 510/50 bandpass emission filter (Chroma) and was captured by a scientific CMOS camera (Hamamatsu Orca Flash 4.0). Custom LabView (National Instruments) software controlled the microscope as previously described^85,86^.

Violet light visual stimulus was performed through a 10x objective (Olympus XLPLN10XSVMP, NA 1.0) by aligning a 405 nm laser (Coherent Obis LX 1284371, 100mW) out of the primary field of view to minimize optical crosstalk into the GFP channel. Laser power was set to 20 mW. Intersegmental vessels immediately posterior to the swim bladder of 5 dpf larvae were selected for Ca^2+^ imaging.

### Pharmacological perturbations

To inhibit fish muscle contraction, para-aminoblebbistatin dissolved in DMSO (final concentration 0.5%) was diluted to a final concentration of 50 μM in E3 media. Fish mounted in agarose were soaked for >30 minutes at 28.5 °C prior to imaging. To block the neuro-muscular junction, tubocurarine hydrochloride pentahydrate (Sigma Aldrich T2379-100MG) at 2.2 mM was applied to fish for > 1 h at 28.5 °C prior to imaging. To block mechano-sensitive ion channels, Ruthenium red (Sigma-Aldrich R2751-1G) at 50 μM was applied to fish for 8 h at 28.5 ° C prior to imaging. After the study involving ruthenium red treatment, fish were released from their agarose mold and drug was washed out. The following day the fish were observed for swimming behavior and body motion-induced EC Ca^2+^ events to confirm absence of long-term effects of the drug treatment.

### Electrical stimulation of fish motion

A chlorided silver wire was inserted inside a 6 MΩ borosilicate glass pipette (VWR Inc.), filled with E3 media. The pipette was positioned near the fish head. A counter-electrode of chlorided silver was placed near the fish tail. Electrical stimulation was applied using a high-voltage amplifier (Krohn-hite 7600M) to amplify 100-µs square pulse generated by the DAQ card (NI BNC-2090A) to 30-45 V. Electrical stimulation did not affect fish heart-rate, and fish remained viable for several days after electrical stimulation.

### CRISPR/Cas9 knockouts

Cas9 protein (CP01) was acquired from PNA Bio Inc. Sequences of all gene-specific sgRNAs were designed using http://chopchop.cbu.uib.no/^87^. The selected sgRNAs were synthesized following established protocols^88,89^. All Cas9 proteins and home-synthesized sgRNAs were purified, aliquoted and stored at -80 °C until use.

For knockout experiments, a mixture of selected sgRNAs (1 μL of each sgRNA at 300 ng/μL), Cas9 protein (1.2 μL at 2 mg/mL), and phenol red dye (0.3 μL) were freshly prepared 30 minutes prior to injection. Zebrafish zygotes were injected at single-cell stage with 500 pL of mixture through the chorion. At 24-30 hpf, injected zygotes were inspected under a Leica stereomicroscope.

To identify knockout efficiency, genomic DNA of injected and control intact zygotes (10 embryos each group) at desired development stages were extracted using the HotSHOT method^90^. The gene sequences and sgRNA target sites of interest were amplified by Platinum Tag PCR reaction (Life Technologies 11304011). The cleavage patterns were visualized by gel electrophoresis using 2% agarose gel, 1x TAE buffer (Fisher Scientific AM9869), 6x purple loading dye (NEB B7024S), and TriDye 100 bp DNA ladder (NEB N3271S). For sequence analysis of cleaved gene sites, the sgRNA target sites were cloned into pENTR/D-TOPO vector (Thermo Fisher K240020), and sent for Sanger sequencing (Genewiz, Inc.). The mutant sequences were identified by comparison to wild-type sequence from intact zygotes.

To knock out *piezo1* expression, we designed five sgRNAs that mutated the 5’ untranslated region (5’-UTR) of the *piezo1* locus, and another five sgRNAs that mutated the start region of first exon, including start codon (Fig. S5A). We confirmed this 351-base pair excision by gel electrophoresis (Fig. S5B) and sequencing (Fig. S5C) of CRISPR/Cas9 injected embryos. To knock out *pkd2* expression, we designed 2 groups of sgRNAs to target two regions in the first exon, separated by 340 base pairs (Fig. S5A). We confirmed this 340-base pair excision by gel electrophoresis (Fig. S5B) and sequencing (Fig. S5C) of CRISPR/Cas9 injected embryos. The TRPV4 gene of zebrafish has relatively short Exon 1 and Exon 2, yielding limited sites for sgRNA binding. We thus designed 2 groups of sgRNAs to mutate 2 regions within its long Exon 3, separated by 560 base pairs (Fig. S5A). We confirmed this 560-base pair excision by gel electrophoresis (Fig. S5B) and sequencing (Fig. S5C) of CRISPR/Cas9 injected embryos. All guide RNA sequences are included in Supplementary File 1. All guide RNA sequences are provided in Supplementary File 1. All genotyping primers are provided in Supplementary File 2.

### Morpholino/Photo-morpholino knockdown

All antisense morpholino oligonucleotides and sense photo-cleavable morpholino oligonucleotides were purchased from Gene Tools, LLC. The concentration of morpholino in solution was determined using a spectrophotometer with a 1 cm path length (Thermo Scientific NanoDrop 2000/2000c). The light-induced cleavage of photo-morpholino was verified by mass spectroscopy at Gene Tools, LLC. For the Piezo1 knockdown experiments by photo-morpholino, each embryo of *Tg (kdrl:Gal4:UAS:GCaMP5G)* or control wild-type (AB or PE) line at single-cell stage was injected with 1.16 nL of duplex, containing 2.5 ng of 1.016 mM Piezo1 antisense morpholino oligonucleotides and 2.5 ng of 1.029 mM Piezo1 sense photo-cleavable morpholino oligonucleotides. At 3 dpf, the injected and control embryos were exposed to 355 nm laser for 30 seconds to cleave the photo-sensitive antisense morpholino. At 5 dpf, functional calcium imaging of larvae were carried out on a custom-built epi-fluorescence microscope, followed by lysis treatment for western blotting analysis.

### Immunocytochemistry and staining

Fish were fixed in fresh 4% EM-grade PFA in PBS (Thermo Fisher AAJ61899-AK) and 0.25% Triton (Sigma Aldrich X100) in PBT (PBS, 0.2% bovine serum albumin (BSA), and Tween-20 0.1%) overnight at room temperature, followed by 3 rinses for 5-min each in PBT. To permeabilize the fish, the samples were incubated in 0.05% Trypsin-EDTA solution on ice for 1 h, followed by 3 rinses for 10-min each in PBT. The samples were then blocked in mixture of 2% Normal Goat Serum, 1% BSA, and 1% DMSO in PBT for 1 h at room temperature. Primary Tie-2 antibody (R&D system, AF928) was diluted 1:1000 in 1% BSA and 1% DMSO in PBT. Samples were incubated in primary antibody solution overnight at 4 °C, followed by 3 rinses for 15-min each in PBT at room temperature. Samples were then incubated in secondary antibody solution overnight at 4 °C, followed by 3 rinses of 15-min each in PBT at room temperature. Samples were manually mounted in low-melting-point agarose for imaging and measurement under laser scanning confocal fluorescence microscopy. Samples were stored in PBS 0.02% sodium azide at 4 °C.

### Western blot analysis

After calcium imaging measurements, all recorded and control fish were separately lysed in ice-cold RIPA lysis buffer (Thermo Fisher Scientific R0278) complemented with 1:100 protease inhibitor (Sigma Aldrich 11697498001). To quantify and compare their Piezo1 expression, β-tubulin expression was used as loading control. Following incubation at 4 °C for 20 minutes, the lysed solution was centrifuged at 16,000 g for 1 hour at 4 °C, and supernatant was retained, followed by the protein concentration determination by microBCA protein assay at 562 nm (Thermo Fisher Scientific 23235). For single fish at 6 dpf, we generally get 100 uL of lysed solution at 0.5 mg/mL protein concentration. Each lysed solution were added with 95 °C 5x Laemmli buffer (2.5 mL of 1.25 M Tris at pH 6.8, 5 mL of 100% glycerol, 1g of electrophoresis grade SDS, 0.8 g of electrophoresis grade DTT, and 1 mL of 1% bromophenol blue) and sonicated briefly for 30 seconds. The protein solutions were then boiled at 37 °C for 10 minutes, followed by brief vortex for 30 seconds and spin down (14,600 rcf, Relative Centrifuge Force) at room temperature.

The protein solutions were loaded into NuPAGE Novex 3-8% Tris-Acetate gels (Thermo Fisher Scientific EA0375BOX) with ∼40 μg protein mass per lane and incubated in NuPAGE Tris-Acetate SDS running buffer (Thermo Fisher Scientific LA0041). Since Piezo1 protein has large molecular weight ∼270 kDa^42^, a Spectra multicolor high range protein ladder (Thermo Fisher Scientific 26625) was chosen for reference. The electrophoresis was carried out for 20 minutes at 80 V followed by 1 hour at 120 V. Gels and 2 pieces of fresh pre-cut extra-thick blotting paper (Bio-rad 1703965) were rinsed in cold 4 °C NuPAGE transfer buffer (Thermo Fisher Scientific NP0006-1) for 2 minutes. Prior to blotting, an immune-blot PVDF membrane (Bio-rad 4569035) was activated in 100 % MeOH for 1 minute and rinsed in same 4 °C NuPAGE transfer buffer for 2 minutes. The transfer sandwich was assembled in the order: extra-thick blotting paper, activated PVDF membrane, gel, extra-thick blotting paper. Air bubbles trapped inside were squeezed out by a roller. A Bio-rad semi-dry and rapid blotting system was used to complete the transfer process at 2.5 A for 15 minutes.

Following transfer, the blotted membranes were inspected under light and rinsed in casein blocking buffer (Sigma Aldrich B6429) overnight at 4 °C. Piezo1 protein is a trimer, containing ∼2,547 AA (∼900 kDa), and becomes dissociated into monomer under denaturing conditions. Membranes were split along 150 kDa mass line into 2 pieces, which contained the Piezo1 monomer (∼270-300 kDa)^42^ and β-tubulin (55 kDa; loading control), respectively. The split membranes were incubated with 10 mL of primary Piezo1 rabbit polyclonal antibody (1: 5000 in Casein blocking buffer, Fisher Scientific 15939-1-AP) and primary β-tubulin rabbit polyclonal antibody (1: 5000 in Casein blocking buffer, Abcam ab6046) at room temperature for 1 hour, followed by 5 rinses of 10 min each in TBST buffer at pH 7.62 (20 mM Tris, 150 mM NaCl, and 0.05% Tween 20). All rinsed membranes were incubated with peroxidase goat anti-rabbit IgG secondary antibody (1:2000 in Casein blocking buffer, Jackson Immuno Research Inc. 111-036-045) at room temperature for > 2 hours, followed by 5 rinses of 10 min each in same TBST buffer. The rinsed membranes were added with 1 mL of chemiluminescence solution mixed from GE Healthcare Amersham ECL Prime western blotting detection reagent (Fisher Scientific RPN2232) and imaged immediately by an Azure Biosystems C400 imager at room temperature. The intensity of protein bands were quantified with normalization by loading control, β-tubulin, and compared by ImageJ and Adobe Photoshop.

### Data analysis and statistics

Data analysis, statistics, and plotting were performed in MATLAB and Python (using Numpy, Scipy, and related libraries). Relative motion was determined for whole-fish recordings by selecting a patch with features of high spatial frequency and measuring displacement by spatial cross-correlation of this patch. For visual stimulus experiments where displacement was large relative to the field of view, relative motion was instead determined using the spatial Fourier transform of a strip of video along the animal body axis. The amount of low frequency signal in the power spectrum, or “blurring” beyond the time resolution of the recording, relative to that at rest was used as an indication of motion. Motion correction was performed using least-squares estimation of Euclidean transformation matrices between frames at a manually adjusted spacing (MATLAB “imregtform”). These transformation matrices were then linearly interpolated between frames. Segmentation of individual vessels was performed manually for whole fish recordings and using Segment Anything 2 (a deep learning model) for the visual stimulus experiment^91^. Analysis of Ca^2+^ imaging data was then performed as previously described^67^.

## Acknowledgement

This work was supported by funds from Harvard University and the Howard Hughes Medical Institute (HHMI). We thank Calum A. MacRae for gift of *Tg (kdrl:Gal4)* line. We thank Jennifer Hou, Christopher A. Werley, Peng Zou, James Lee, Erin Yue Song, Eva Fast, Joel M. Kralj, and Adam Douglass for helpful discussions and training. We gratefully acknowledge help from members of the labs of Adam Cohen, Florian Engert, Alex Schier and Leonard Zon. Florian Engert received funding from the National Institutes of Health (U19NS104653 and 1R01NS124017-01), DoD (W911NF2420112) and the Simons Foundation (SCGB 542973 and NC-GB-CULM-00003241-02). This work was supported by funding from the National Heart, Lung, and Blood Institute (1P01HL131477) and the National Institute of Diabetes and Digestive and Kidney Diseases (R01 DK140372) to L.I.Z.

