## Supplementary Information for "Swimming motions evoke Ca^2+^ events in vascular endothelial cells of larval zebrafish via mechanical activation of Piezo1"

### Contents

Supplementary Table 1

Supplementary Figures 1-5

Supplementary Video Captions 1 - 4

| Modality | Stress | Strain | Timescale |
|---|---|---|---|
| (a) Shear | 10 dyn/cm$^2$ = 1 N/m$^2$ | NA | ~0.25-1 s |
| (b) Pressure | 22.6~66.5 N/m$^2$ | 1.6~4.1% | ~0.25-1 s |
| (c) Circumferential stretch (vascular smooth muscle) | 0~4,000 N/m$^2$ | 10-20 % | ~1-10 s |
| (d) Axial stretch (skeletal muscle) | 50,000~69,000 N/m$^2$ | 2~6 % | ~0.01-0.1 s |

**Supplementary Table 1. Approximate mechanical stress, strain, and timescale of forces on ECs *in vivo*. (a)** Blood shear stress τ is derived from $\tau = \gamma \frac{dv}{dr}$, where γ (0.01 Pa s) is the blood viscosity, and dv/dr is the flow velocity gradient at the vessel wall. For embryonic zebrafish at 5 dpf, the radius of the dorsal aorta (DA) is *R* ~10 μm, maximum centerline blood velocity is 1000 - 2000 μm/s, and dv/dr is estimated as 100 - 200 $s^{-1}$ [85]. Shear stress τ is estimated as 1 - 2 Pa. **(b)** Pulsatile blood pressure of embryonic zebrafish (systolic – diastolic pressure) varies with age. For larval zebrafish (84-96 hpf), pulsatile blood pressure in arteries is 0.2 - 0.5 mmHg (26 - 67 Pa)[86], while for adult zebrafish (3 months), pulsatile pressure in dorsa aorta (DA) is 1 - 1.5 mmHg (133 - 200 Pa)[87]. The circumferential strain (dR/R) on dorsal aorta is estimated by: $\frac{dR}{R} = \frac{q\,R}{E\,t}$, where *R* is the radius of blood vessel (~10 μm), *q* is the pulsatile blood pressure (26 - 67 Pa), *E* is the elastic modulus of ECs (~4 kPa), and *t* is the thickness of vessel EC wall (~4 μm)[88]. The strain is estimated to be ~1.7 - 4.1%. Timescales for (a) and (b) are based on the duration of a single heartbeat at 5 dpf[89]. **(c)** Traction force of smooth muscle cell measured *in vitro* is ~0 – 4 kPa[90]. The circumferential strain is derived from vasoconstrictor treatments in larval zebrafish[91,92]. The timescale is derived from prior measurements of spontaneous contraction or phenylephrine-induced contraction on arterial explants[93]. **(d)** The axial stress caused by skeletal muscle cells and its timescale are derived from literature[42]. The axial strain is estimated by our *in vivo* whole body fluorescence imaging.

**Supplementary Figures**

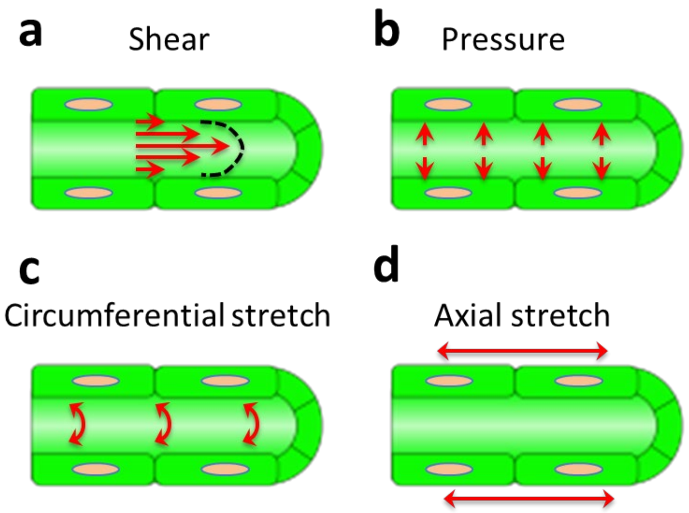


Fig S1: ECs experience diverse forces *in vivo*. **(a)** Pulsatile blood flow applies shear forces along the inner wall of vessels. **(b)** Pulsatile blood pressure produces an outwards push and causes radial stretch on ECs. **(c)** Smooth muscle cells or mural cells (both distributed sparsely on vessels of embryonic zebrafish[54]) slowly contract the vessels and cause circumferential stretch. **(d)** Body skeletal muscles rapidly contract to generate swimming motions, applying longitudinal tension and leading to axial stretch.

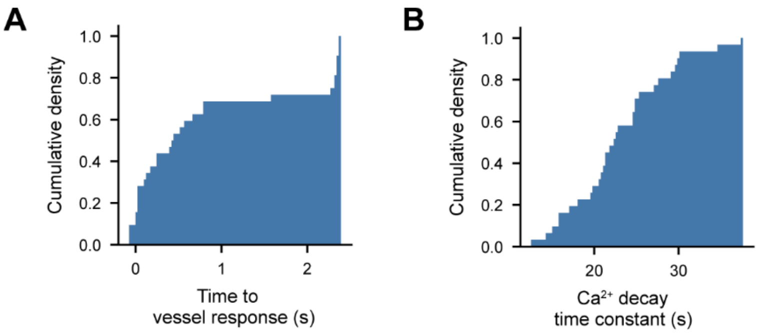


Fig. S2: Temporal structure of motion-triggered endothelial cell $Ca^{2+}$ dynamics. (A) Cumulative distribution of time-delay between onset of twitch motion and EC GCaMP6s to reach 10% of its peak intensity increase. (B) Time constant of EC GCaMP6s decay based on mono-exponential fit. 31 violet light-triggered $Ca^{2+}$ events from intersegmental vessels in *n = 2 Tg(kdrl:Gal4; UAS:GCaMP6s)* fish at 5 days post fertilization.

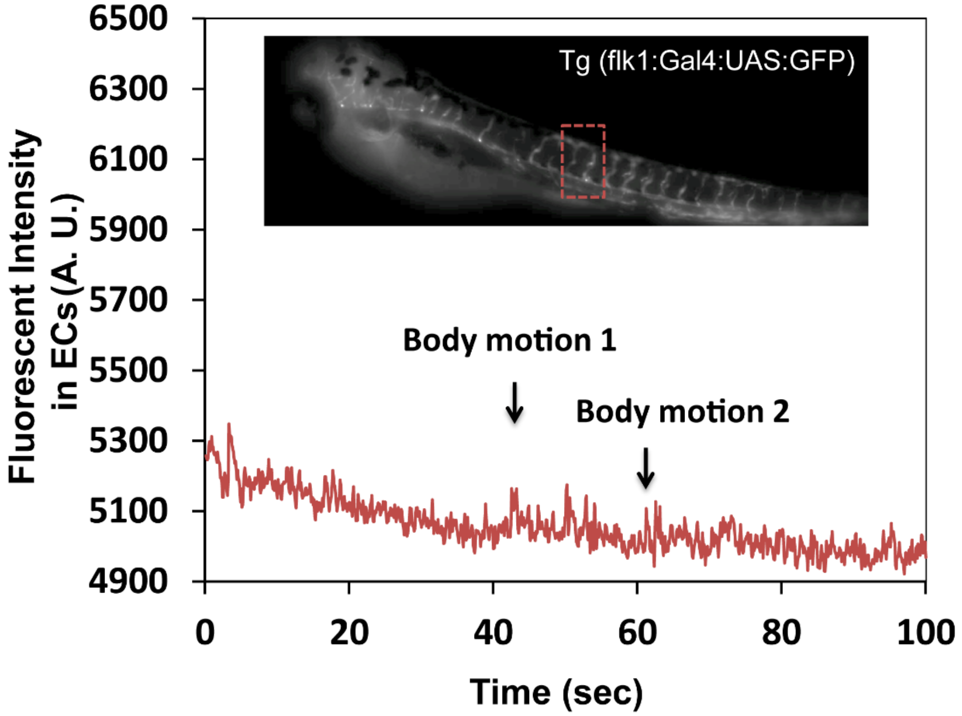


Fig. S3: Motion-induced signal in *Tg(kdrl:Gal4; UAS:eGFP)* fish**.** Motion-induced fluorescence transients in ECs expressing eGFP were minimal and sharply peaked. This confirms that the signals observed in fish expressing $Ca^{2+}$ sensors were due to changes in intracellular $Ca^{2+}$ level, and not due to motion artifacts. Inset: fluorescent image of zebrafish, *Tg(kdrl:Gal4; UAS:eGFP)*, 5 days post fertilization.

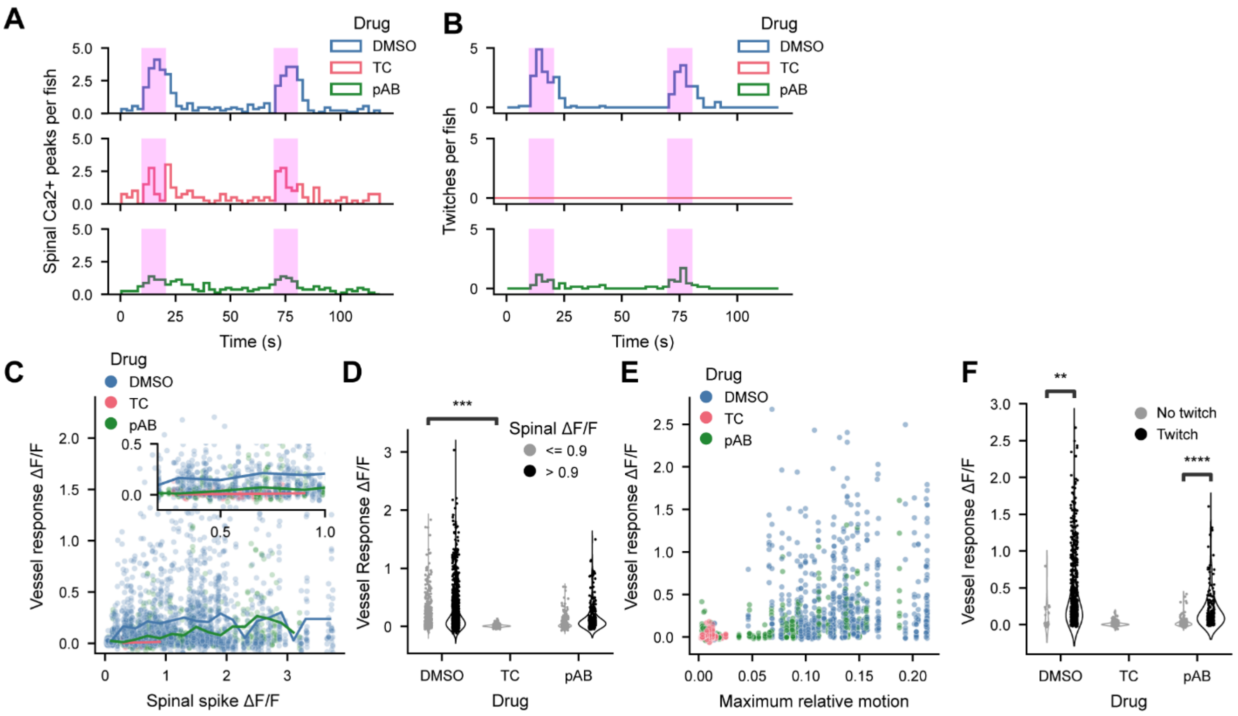


Fig. S4: Dynamics of violet light-evoked $Ca^{2+}$ responses in *Tg(kdrl:Gal4; vglut2a:Gal4; UAS:GCaMP6s)* fish. (A) Spinal $Ca^{2+}$ spikes binned by time (TC = tubocurarine 2.2 mM, pAB = para-amino blebbistatin 50 μM) over all trials and fish in Figure 3. Violet light stimuli are indicated by pink bars. (B) Body twitches binned by time. (C) Individual vessel segment $Ca^{2+}$ responses triggered on spinal $Ca^{2+}$ spikes (DMSO: 2459 vessel segment events, TC: 545 vessel segment events, pAB: 750 vessel segment events). (D) Data from (C) binned based on a spinal $Ca^{2+}$ ΔF/F threshold of 0.9. (E) Individual vessel segment $Ca^{2+}$ responses triggered on peaks in relative motion (DMSO: 680 events, TC: 320 events, pAB: 448 events). (F) Data from (E) binned based on a relative motion threshold of 0.068. $n$ = 9 fish DMSO, $n$ = 4 fish TC,$n$ = 8 fish pAB in all panels. Statistical tests: mixed-effect linear model considering effects of drug and individual fish on each vessel; (D) TC vs DMSO: $z$ = 3.827, $p$ = 1.30e-4; (F) DMSO: $z$ = 2.95, $p$ = 3.19e-3; pAB: $z$ = 9.60, $p$ = 8.13e-22.

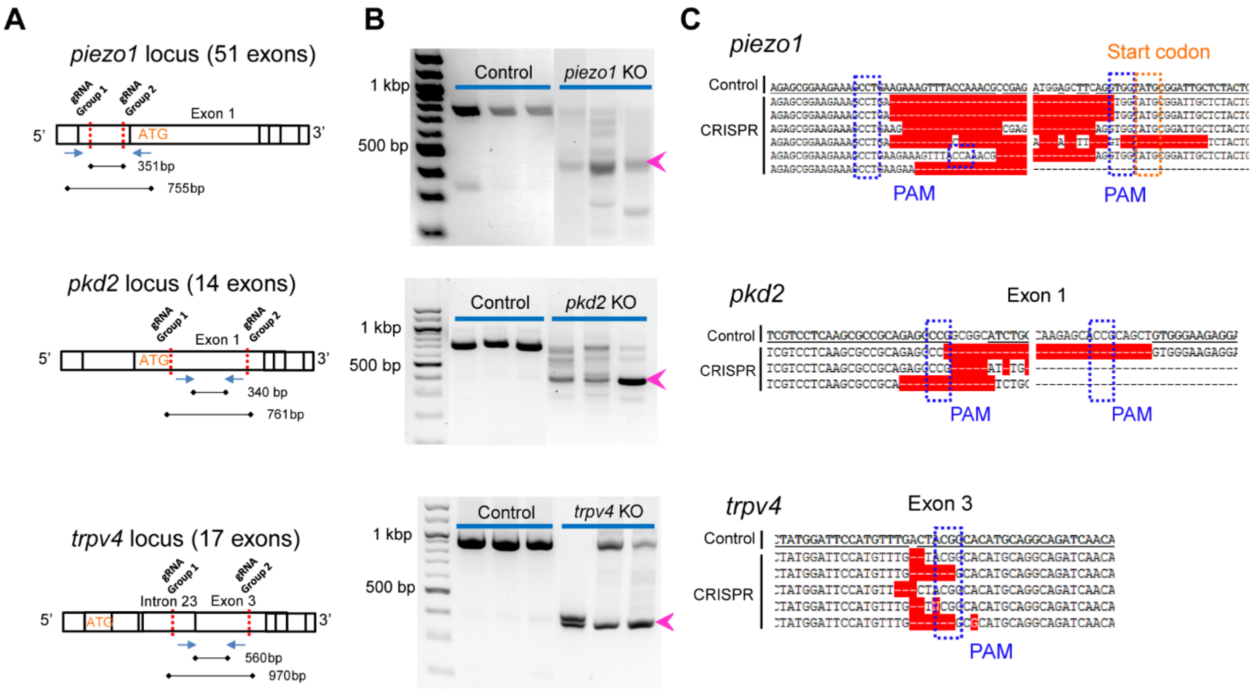


<u>Fig. S5: Design and validation of CRISPR/Cas9 knockout for mechanosensitive ion channels.</u> Top, middle, and bottom rows correspond to *piezo1*, *pkd2*, and *trpv4* respectively. (A) gRNA target design on each genomic locus. Full guide sequences are provided in Supplementary File 1. (B) DNA gel electrophoresis of genomic DNA after knockout performed in *Tg(kdrl:Gal4;UAS:GCaMP5G)* fish. Larvae injected with the target gRNA in each experiment were genotyped after $Ca^{2+}$ imaging. Only the individuals with the correct bands shown here were retained for analysis of $Ca^{2+}$ imaging data. Gel lanes for injected fish with unsuccessful knockout (i.e. wild-type or uninterpretable band) have been cropped out. (C) Sanger sequencing validation of gRNA targeting.

**Supplementary Video Captions**

**Supplementary Video 1: Endothelial $Ca^{2+}$ dynamics evoked by spontaneous motion in a 7-day old *Tg(kdrl:Gal4; UAS:GCaMP5G)* larva.**

**Supplementary Video 2: Violet visual stimulus-evoked motion and $Ca^{2+}$ dynamics under drug treatment in 5-day old *Tg(kdrl:Gal4;vglut2a:Gal4;UAS:GCaMP6s)* larvae.** TC = tubocurarine 2.2 mM, pAB = para-amino blebbistatin 50 µM. Magenta circle indicates 405 nm visual stimulus. Scale bar 25 µm.

**Supplementary Video 3: Visual stimulus-evoked $Ca^{2+}$ transients in the absence of swimming motion at junctions between the dorsal aorta and intersegmental arteries.** 5-day old *Tg(kdrl:Gal4;UAS:GCaMP6s)* larva treated with 2.2 mM tubocurarine. Magenta circle indicates 405 nm visual stimulus. Scale bar 25 µm.

**Supplementary Video 4: Blood flow in pAB-treated and control larvae under violet visual stimulus.** Magenta circle indicates 405 nm visual stimulus. Rectangles demarcate a segment of the dorsal aorta and posterior cardinal vein where blood flow is visible in the control fish. Scale bar 50 µm.
